# Phenotypic characterization of cryptic species in the fungal pathogen *Histoplasma*

**DOI:** 10.1101/2024.01.08.574719

**Authors:** Victoria E. Sepúlveda, Jonathan A. Rader, Jingbaoyi (Janet) Li, William E. Goldman, Daniel R. Matute

## Abstract

Histoplasmosis is an endemic mycosis that often presents as a respiratory infection in immunocompromised patients. Hundreds of thousands of new infections are reported annually around the world. The etiological agent of the disease, *Histoplasma,* is a dimorphic fungus commonly found in the soil where it grows as mycelia. Humans can become infected by *Histoplasma* through inhalation of its spores (conidia) or mycelial particles. The fungi transitions into the yeast phase in the lungs at 37°C. Once in the lungs, yeast cells reside and proliferate inside alveolar macrophages. We have previously described that *Histoplasma* is composed of at least five cryptic species that differ genetically, and assigned new names to the lineages. Here we evaluated multiple phenotypic characteristics of 12 strains from five phylogenetic species of *Histoplasma* to identify phenotypic traits that differentiate between these species: *H. capsulatum sensu stricto*, *H. ohiense*, *H. mississippiense*, *H. suramericanum*, and an African lineage. We report diagnostic traits for two species. The other three species can be identified by a combination of traits. Our results suggest that 1) there are significant phenotypic differences among the cryptic species of *Histoplasma*, and 2) that those differences can be used to positively distinguish those species in a clinical setting and for further study of the evolution of this fungal pathogen.

**IMPORTANCE:** Identifying species boundaries is a critical component of evolutionary biology. Genome sequencing and the use of molecular markers have advanced our understanding of the evolutionary history of fungal pathogens, including *Histoplasma*, and have allowed for the identification of new species. This is especially important in organisms where morphological characteristics cannot be used for this purpose. In this study, we revise the taxonomic status of the four named species of the genus *Histoplasma*: *H. capsulatum sensu stricto*, *H. ohiense*, *H. mississippiense*, and *H. suramericanum* and propose the use of species-specific phenotypic traits to aid their identification when genome sequencing is not available. These results have implications not only for evolutionary study of *Histoplasma*, but also for clinicians, as the *Histoplasma* species could determine the outcome of disease and treatment needed.

## INTRODUCTION

*Histoplasma* is an ascomycete dimorphic fungus and the causal agent of histoplasmosis. This disease is arguably the most common fungal respiratory infection, with hundreds of thousands of new infections occurring annually worldwide (Cano and Hajjeh 2001; Kauffman 2007; Hage *et al*. 2015; Bongomin *et al*. 2019). The disease is particularly common in immunocompromised patients and the majority of cases are reported in patients that have undergone chemotherapy (Adderson 2004; Hess *et al*. 2017), organ transplant (Freifeld *et al*. 2005), or are suffering from AIDS (Adenis *et al*. 2018; Myint *et al*. 2020). Histoplasmosis is not of mandatory report, and for that reason, the true disease burden remains largely unknown (Hage *et al*. 2015; Armstrong *et al*. 2018; Oladele *et al*. 2018; Scully and Baddley 2018; Adenis *et al*. 2018). Nonetheless, there are indications that the disease is more important than is currently understood. Histoplasmin skin reactivity tests suggest that by age 20, more than 90% of individuals residing in the Continental United States are skin test-positive for a previous infection, or at least exposure to the pathogen (Manos *et al*. 1956). Over 100 outbreaks were reported in the twentieth-century in the USA (Benedict and Mody 2016). Similar population assessments indicate that a large proportion of the population has been exposed to *Histoplasma* at some point in their life. Among immunosuppressed patients, the population most at risk, 25% of AIDS patients living in endemic regions of *Histoplasma* develop histoplasmosis; untreated cases usually lead to patient death and infected individuals often need intense and prolonged antifungal therapy (Kauffman 2007; Hage *et al*. 2015).

Producing the sexual stage of *Histoplasma* in laboratory conditions is exceedingly difficult (but see (Kwon-Chung 1972a; b, 1973; Muniz *et al*. 2014)), which has made the study of potential species boundaries challenging for decades. Initial assessments of diversity proposed three different subspecies *for Histoplasma capsulatum. Histoplasma capsulatum* var. *capsulatum* was thought to mainly be found in human patients and caused the classical pulmonary form of histoplasmosis, *H. capsulatum* var. *duboisii* allegedly caused a milder version of the disease with granulomatous lesions in skin and osseous tissues, and *H. capsulatum* var*. farciminosum* was thought to be a pathogen of mules and horses (Ajello 1968). The application of phylogenetics using molecular markers revealed that these lineages were artifactual and did not follow the evolutionary history of the pathogen (Kasuga *et al*. 1999, 2003). Multilocus sequence typing revealed at least six lineages within *Histoplasma* (Kasuga *et al*. 2003). Another classification of *Histoplasma* is based on the presence/absence of the polysaccharide α-(1, 3)-glucan in the cell wall (*AGS1* locus), produced only during the yeast phase. Strains that possess α-(1, 3)-glucan have a rough colony morphology and are classified as chemotype 2 strains, which represent the majority of the strains found worldwide. Strains that lack α-(1, 3)-glucan have a smooth colony morphology, are classified as chemotype 1 strains, and are restricted to a North American lineage (Domer 1971; Reiss 1977; Reiss *et al*. 1977; Rappleye *et al*. 2004; Edwards *et al*. 2011). The virulence requirements for α-(1, 3)-glucan have been shown to differ among *Histoplasma* lineages (Edwards *et al*. 2011). Additional, as-yet unidentified lineages are likely to exist within *Histoplasma* (Kasuga *et al*. 2003; Teixeira *et al*. 2016)

The implementation of genome sequencing confirmed the existence of differentiated genetic lineages and revealed that these clades were sufficiently diverged to be considered phylogenetic species (Sepúlveda *et al*. 2017; Almeida-Silva *et al*. 2021; Jofre *et al*. 2022). Five species satisfied the first assessment of genome concordance and differentiation: *H. ohiense*, *H. mississippiense*, *H. capsulatum sensu stricto*, *H. suramericanum*, and a *Histoplasma* lineage from continental Africa. Additional genome sequencing revealed the existence of two additional phylogenetic species, one endemic to the Indian subcontinent (Jofre *et al*. 2022), and one endemic to Southern Brazil (Almeida-Silva *et al*. 2021). These seven species in the *Histoplasma* genome diverged over 1.5 million years ago and have accrued extensive genetic differences that make them advanced along the speciation continuum (Sepúlveda *et al*. 2017).

The taxonomic rearrangement of the *Histoplasma* genus set the basis for further studies and propelled important developments in understanding the biology of *Histoplasma*. Genome assembly of strains from each of these species suggested genome content differences and rearrangements which, in turn, have suggested a rapid turnover of genome structure in the genus (Voorhies *et al*. 2022). Surveys of gene exchange have also revealed low levels of admixture among lineages which indicates that hybridization might be of importance in the evolution of *Histoplasma* (Maxwell *et al*. 2018; Jofre *et al*. 2022). From a more applied perspective, Sepúlveda et al. (2017) reported extensive genetic differences along the genome and the possibility of using molecular markers for molecular detection which could be harnessed by clinical researchers and inform the epidemiological patterns of each of these lineages.

Despite all the genomic progress, no systematic assessment has been performed to determine whether these phylogenetic species differ phenotypically. Clearly there is extensive genetic differentiation in the genus, but taxonomic revisions should be accompanied by descriptions that can serve clinical and evolutionary researchers alike (Matute and Sepúlveda 2019; Chethana *et al*. 2021). The initial species description suggested that previous assessments of phenotypic differentiation in *Histoplasma* might follow species boundaries (Sepúlveda *et al*. 2017). Nonetheless, no survey has measured potential intraspecific and interspecific variation in common conditions. Here we bridge that gap. We explored whether the genetic differentiation within *Histoplasma* might explain some variability in the group and whether phenotypic variation follows species boundaries. In this report, we quantified four phenotypic traits and found yeast-culture-based diagnostic characters for two of the *Histoplasma* species, *H. ohiense*, and *H. mississippiense*. The other three species, *H. capsulatum, H. suramericanum*, and the African lineage, can be identified by a combination of multiple traits. Using this information, we revise the taxonomic status of the four named species of the genus *Histoplasma*.

## MATERIALS AND METHODS

### Fungal strains and culture conditions

*Histoplasma* isolates used in this study were donated to W.G. during a span of 15 years. Information pertinent to each isolate is listed in Table 1. All isolates were kept in 15% glycerol at -80°C until they were ready to be subcultured. An aliquot of the frozen culture was streaked into *Histoplasma* Macrophage Medium (HMM) plates. Strains were then grown in HMM (solid or liquid) at 37°C with 5% CO2 as previously described ^(^Worsham and Goldman 1988^)^. Solid medium contained 0.6% agarose (SeaKem ME grade) and 25 mM FeSO4. All liquid cultures were incubated at 37°C with 5% CO_2_ on an orbital shaker (Infors HT Multitron) at 150 rpm. All reference strains were deposited in the Westerdijk Fungal Biodiversity Institute CBS collection (Table 1).

**TABLE 1.** *Histoplasma* Isolates used in this study.

| Isolate | Phylogenetic species *sensu* Kasuga et al. (2003) | Phylogenetic species *sensu* Sepúlveda et al. (2017) | Origin | Year collected | Other Designations |
| --- | --- | --- | --- | --- | --- |
| G186A | H81 lineage | *H. capsulatum* | Panama | 1967 or before | ATCC 26029<br>CBS145494 |
| G184A | H81 lineage | *H. capsulatum* | Panama | 1967 or before | ATCC 26027 |
| G217B | NAm 2 | *H. ohiense* | Louisiana/USA<br>A | 1973 or before | ATCC 26032<br>CBS145495 |
| CI_17 | NAm 2 | *H. ohiense* | Missouri/USA | Unknown | CBS145496 |
| WU24 | NAm 1 | *H. mississippiense* | Missouri/USA | 2004 | CBS145497 |
| CI_19 | NAm 1 | *H. mississippiense* | Missouri/USA | 2003 | CBS145498 |
| 3_11G | N/A | *H. suramericanum* | Guatemala | 2011 | CBS145499 |
| 21_14 | N/A | *H. suramericanum* | Guatemala | 2014 | N/A |
| 27_14 | N/A | *H. suramericanum* | Guatemala | 2014 | N/A |
| *H. duboisii* | Africa | *H. capsulatum* var. *duboisii* | ? | Unknown | N/A |
| H88 | Africa | N/A | Belgium,<br>probably from<br>a former<br>Belgian colony | 1975 or before | ATCC 32281<br>RV26821 |
| H143 | Africa | N/A | South Africa | 1954 | CBS 287.54 |

### Yeast colony morphology

We scored the yeast colony morphology of 12 *Histoplasma* isolates (at least two isolates from each species). For each isolate, we added 10 **μ**l of a late exponential phase culture on a HMM plate. We grew 36 aliquots per Petri dish. We incubated plates at 37°C in 5% CO_2_ for at least 10 days before we imaged each colony. Colonies were classified as rough or smooth. To ensure reproducibility, we scored at least 12 colonies per species but no species showed intralineage variation in colony morphology. To compare the proportions of rough vs. smooth colonies among species, we used a 2-sample test for equality of proportions with continuity correction (function *prop.test*, library *stats*, R Core Team 2018).

### Evaluation of extracellular proteolytic activity

The second trait we evaluated was proteolytic activity. Several studies have reported the existence of extracellularly secreted serine proteases in *Histoplasma*. In particular, isolates from the RFLP1 group (later named *H. mississippiense*) were the only ones that manifested this phenotype (Zarnowski *et al*. 2007 *cf*. Muotoe-Okafor *et al*. 1996 for reports of proteolytic activity in African strains). To evaluate extracellular proteolytic activity in different species of *Histoplasma*, we grew the 12 *Histoplasma* isolates (Table 1) in HMM plates supplemented with 1.5% skim milk. Strains with a proteolytic activity show a clear halo around their yeast colonies. 15 g of instant nonfat dry milk (Hoosier Hill Farm brand,Middleton, WI ) were reconstituted in 500 ml of distilled water. Once the skim milk was fully dissolved, 6 g of agarose (SeaKem ME grade) were added and autoclaved to make HMM plates as previously described (Worsham PL and Goldman WE, 1988). 10 **μ**l of a late exponential phase culture were spotted onto HMM plates supplemented with skim milk. We spotted 4 strains per plate to allow for any transparent clearance area around fungal spots to appear, indicative of proteolytic activity. We incubated the experiment using the same conditions as described immediately above to study yeast colony morphology. We scored 12 colonies per isolate for presence / absence of a halo, and when present, measured halo size. The size of the halo was measured as the distance from the edge of the colony to the outer edge of the cleared ring. To compare halo sizes, we used a Welch’s Two Sample *t*-test (function *t.test*, library *stats*, (R Core Team 2018)).

### Optical density and growth curves of *Histoplasma* yeast cultures

We also measured the growth rate of different *Histoplasma* genotypes in liquid media. For the growth curves, we inoculated 30 ml of HMM broth with 1 × 10^6^ yeast/ml and grew the culture for 11 days. We repeated this procedure for each of the 12 strains. We removed 600 **μ**l from each culture and mixed them with 300 **μ**l of 3M NaOH in a plastic cuvette, which was covered with Parafilm and vortexed for 10 seconds to separate yeast clumps and measure optical density (OD) in a GENESYS 10vis spectrophotometer (Thermo Spectronic) starting at day 0, and at every 24 hours after that until day 11. To quantify the rate of growth, we used a four-parameter logistic model with the form:

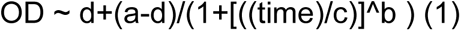

where *a* is the OD at the beginning of the experiment (presumably close to zero), *b* is the rate of increase in OD at point *c*, the inflection point of the curve, and *d* is the maximum OD in the curve, the asymptote. This model allows for an initial growth where cells are dividing but do not increase the OD value and includes an asymptote, calculated from the data, at which cells do not replicate anymore. Since nonlinear logistic regression has difficulties optimizing the values for each of the four constants in the equations, we tried 10 starting values per constant and found the model with the lowest Akaike Information Criterion (AIC, (Akaike 1973)) with the function ‘*AIC’*, (library ‘*stats’*, (R Core Team 2018)). To fit the regressions, we pooled isolates within phylogenetic species.

To determine whether the four fitted parameters differed among species, we generated 1,000 bootstrapped regressions using the R function *nls.boot* (library *nlstools*, (Baty *et al*. 2013, 2015)). We then compared the values of *b*, *c* and *d* across species using non-parametric tests (Wilcoxon rank sum test with continuity correction, function *wilcox.test*, library *stats*, (R Core Team 2018)).

### Yeast Area

Finally, we studied the area of individual yeast cells in different *Histoplasma* isolates. We grew one isolate per *Histoplasma* species to evaluate phenotypic variability between species. Table 1 lists the isolates included in this study. 10 **μ**l from a yeast culture that had large yeast clumps removed were mixed with 10 **μ**l of Lactophenol Cotton Blue on a glass slide. Differential interference contrast (DIC) images were obtained using 100X/1.4 Oil UPlan S Apo PSF quality objective on an Olympus BX-61 microscope and collected using a QImaging RETIGA 4000R color camera and Volocity 6.3 acquisition software. Exposure was adjusted to ensure pixel intensities were not saturated (Pixel size: 0.0608 **μ**m / pixel). Yeast cells were measured by drawing an ellipse around each imaged cell in imageJ (Schneider *et al*. 2012). Ellipse area (in **μ**m^2^) was taken as a measure of cell size. To compare the yeast cell size across different species, we used a linear model in which cell area was the response and the species identity was the grouping factor. We used the R function *aov* (library *stats*, (R Core Team 2018)). Finally, we compared among lineages (all pairwise comparisons) using Tukey contrasts with the R function *TukeyHSD* (library stats, (R Core Team 2018)).

## RESULTS

### *Histoplasma ohiense* differs in their yeast colony morphology

Multiple previous studies have reported variation in yeast colony morphology across isolates of *Histoplasma* (Domer 1971; Reiss 1977; Reiss *et al*. 1977). Some isolates show smooth colonies while some others show rough ones. We studied whether this phenotypic variation was species-specific or whether there was intraspecific variation within five phylogenetic species of *Histoplasma*. Figure 1 shows the yeast colony morphology for 12 different *Histoplasma* isolates after growing at 37°C for 10 days in HMM media. All isolates from four *Histoplasma* species, *H. capsulatum*, *H. mississippiense*, *H. suramericanum*, and the Africa clade, had rough yeast colonies (*n*=12 per species). On the other hand, all the replicates across isolates of *H. ohiense* (*n*=24) had smooth yeast colonies. Not surprisingly, these proportions are significantly different (2-sample test for equality of proportions with continuity correction: *χ*^2^ = 44.083, df = 1, *p* = 3.147×10^-11^). This morphological difference within *Histoplasma* has been attributed to the lack of α-(1,3) glucan in their cell wall (Klimpel and Goldman 1987, 1988; Rappleye *et al*. 2004; Sepúlveda *et al*. 2014). These comparisons indicate that yeast colony morphology is a diagnostic trait of *H. ohiense* and is sufficient to differentiate the species from the other four lineages.

**FIGURE 1.**
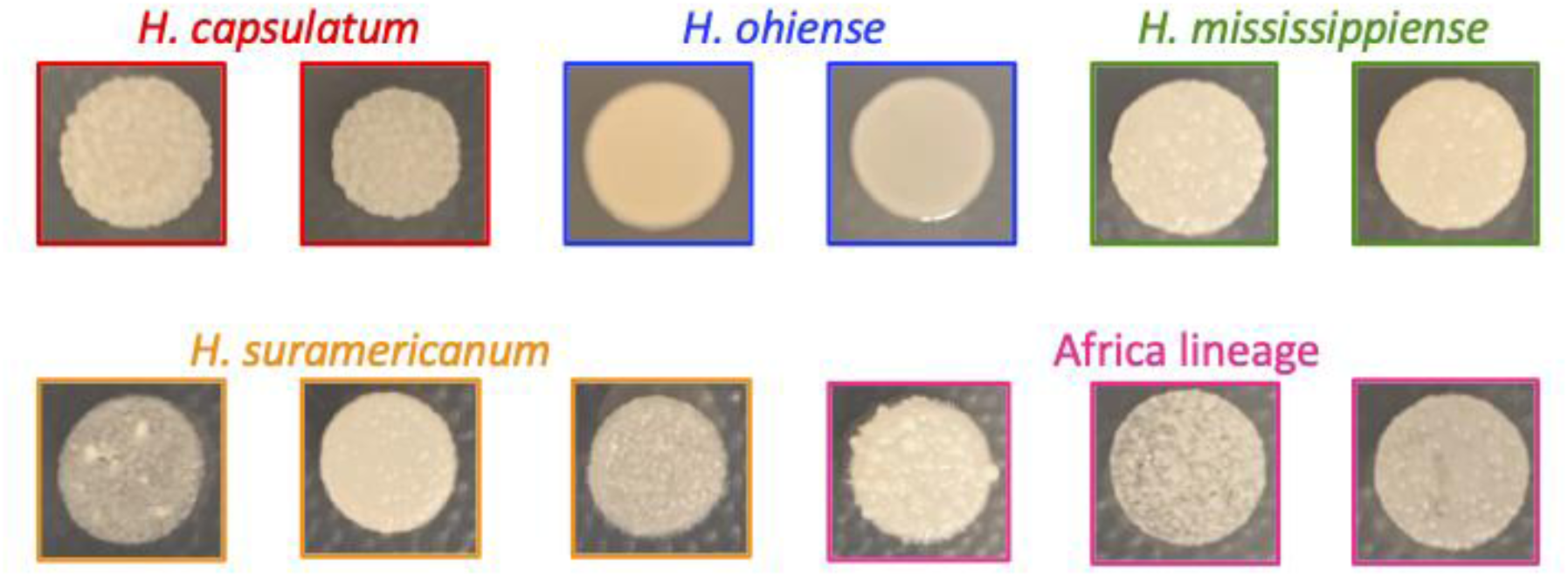
Colony morphology in *Histoplasma* species. Fungal strains were grown on agarose-solidified HMM plates. Smooth morphology occurs in the absence of α-(1,3)-glucan in the cell walls. *Histoplasma ohiense* is the only species that shows a smooth colony morphology due to the lack of α-(1,3)-glucan.

### Production of extracellular proteolytic activity is restricted to *H. mississippiense*

Production of extracellular proteolytic activity using HHM media supplemented with skim milk had been previously described in some isolates of *Histoplasma* (Zarnowski *et al*. 2007). We studied whether the five different species differed in their proteolytic ability. We grew five of the previously identified lineages in skim milk media to determine whether they showed proteolytic activity. Figure 2 shows an example of each of the five species growing as yeast in HMM media supplemented with skim milk at 37°C. Of the five species, *H. mississippiense* was the only lineage to show a clearance halo which is a proxy of the ability of the colony to break down proteins. Within *H. mississippiense*, there was variation in the size of the halo. WU24 had a halo distance of 0.395 cm (SD=0.028), while CI-19 had a halo distance of 0.195cm (SD=0.050) which differed significantly from each other (Welch’s Two Sample *t*-test, *t* = -11.047, df = 14.302, *p* = 2.164×10^-8^) suggesting the existence of intraspecific variation in the genetic mechanisms involved in this trait within *H. mississippiense*, though a halo is always present. The observation is consistent with previous studies that suggested that isolates from this lineage (originally termed RFLP1) are the only ones with a extracellular protease ability (Zarnowski *et al*. 2007), and suggest that proteolytic activity is a diagnostic trait of *H. mississippiense*.

**FIGURE 2.**
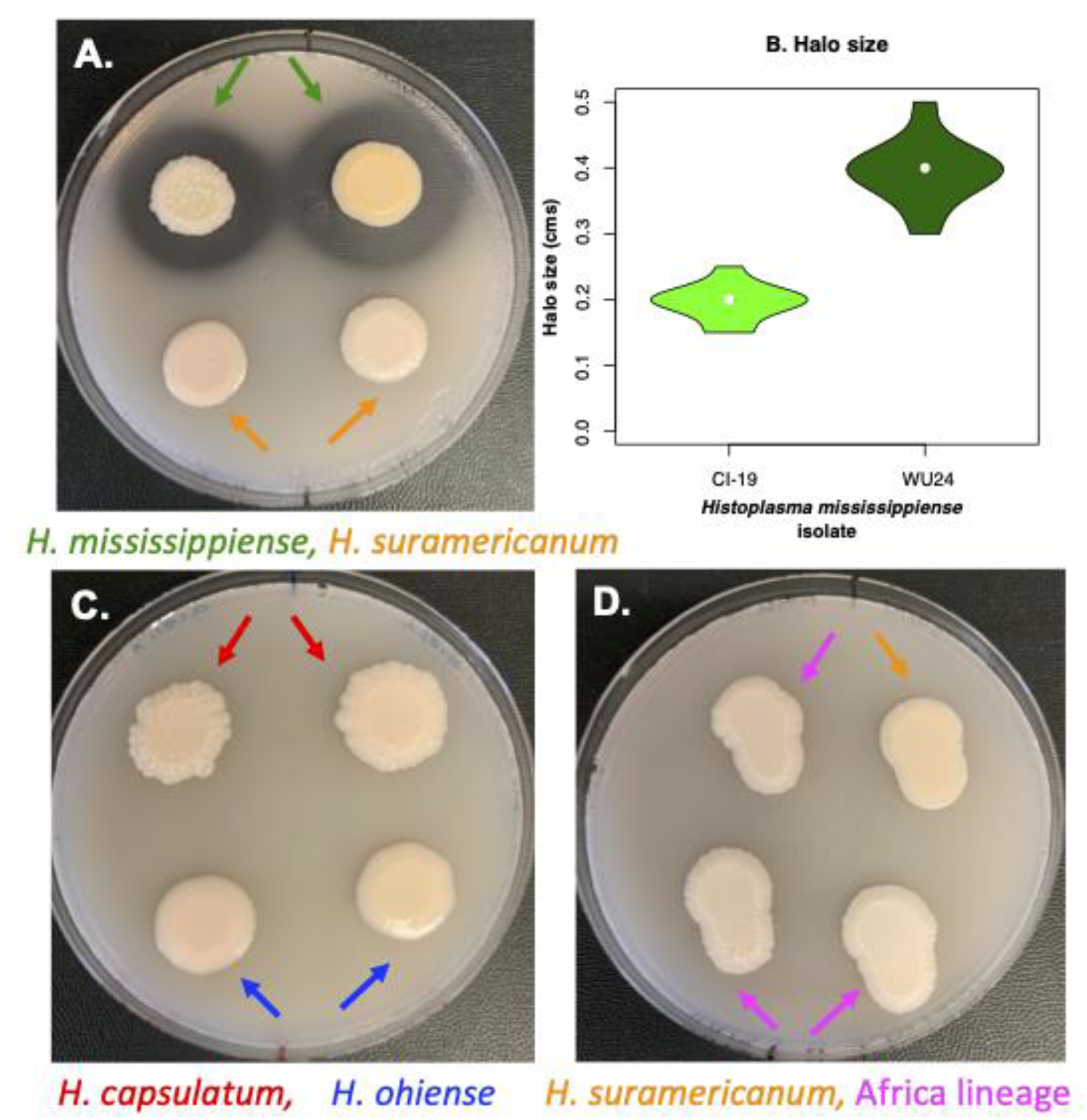
Demonstration of extracellular proteolytic activity in *Histoplasma* isolates. Fungal strains were grown on agarose-solidified HMM supplemented with 1.5% skim milk. Secreted proteolytic activity is visible as transparent clearance halos around fungal colonies was assessed after 10 days of growth at 37°C in 5% CO_2_. The presence of extracellular proteases was observed only in *H. mississippiense* strains (**A**). The size of the halo varied within *H. mississippiense* (**B**). The other four species of *Histoplasma* included in this report showed no proteolytic activity (i.e., no halo; **C** and **D**).

### Growth curves and optical density

We evaluated whether different genotypes of *Histoplasma* had differences in their growth rate and if such differences corresponded with species boundaries. We used optical density as a proxy for the number of cells in a liquid culture, and fitted logistic models that modeled the rate of increase of the different species. Figure 3 shows the results of the best fit for each species. The growth curves of the five species show a better-fit to a logistic dose-response function than to a linear function (Table S1). Non-linear regressions are highly dependent on the seed values used for the optimizations, so we maximized the fit using AIC values (Table S2).

**FIGURE 3.**
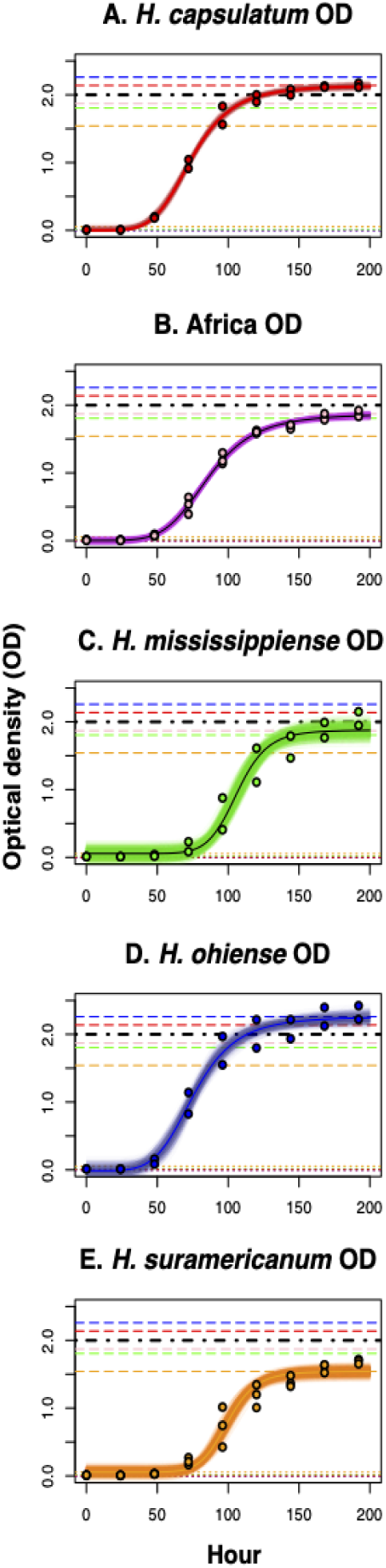
Four-parameter logistic models for the rate of growth of five *Histoplasma* species. All experiments were done in HMM broth media at 37°C with 5% CO_2_. Growth was measured by recording the optical density (OD600) of liquid cultures at different time points (0, 24, 48, 72, 120, 144, 168, and 192). **A.** *Histoplasma capsulatum*. **B.** African lineage **C.** *Histoplasma mississippiense* **D.** *Histoplasma ohiense* **E.** *H. suramericanum*. Semitransparent lines show 1,000 bootstrapped model fits.

We focused on two of the four calculated parameters, the intercept (*a*) and the asymptote (*d*). We found that even though there are significant differences among bootstrapped distributions of the intercept (*a*, Table 2), all intercepts are also centered around zero (Figure 4A). Since the intercepts were similar, comparisons among asymptote (d) values are informative and indicate whether the species have differences in the growth saturation point. Indeed, the values of all the inferred asymptotes differed among the five species, but there were two clearly differentiated groups (Table 2). The growth curves at 196 hours for two of the species (*H. capsulatum* ss, and *H. ohiense*) had OD asymptotes higher than 2. On the other hand, the other three species (*H. mississippiense,* the African lineage, and *H. suramericanum*) had OD asymptotes lower than 2 (Figure 4B, Table 2). This difference can be used to discriminate between these two clusters of species, and suggest that OD-based growth curves can be a taxonomic trait that can aid species identification in *Histoplasma*, but one that does not serve as a diagnostic trait in isolation.

**FIGURE 4.**
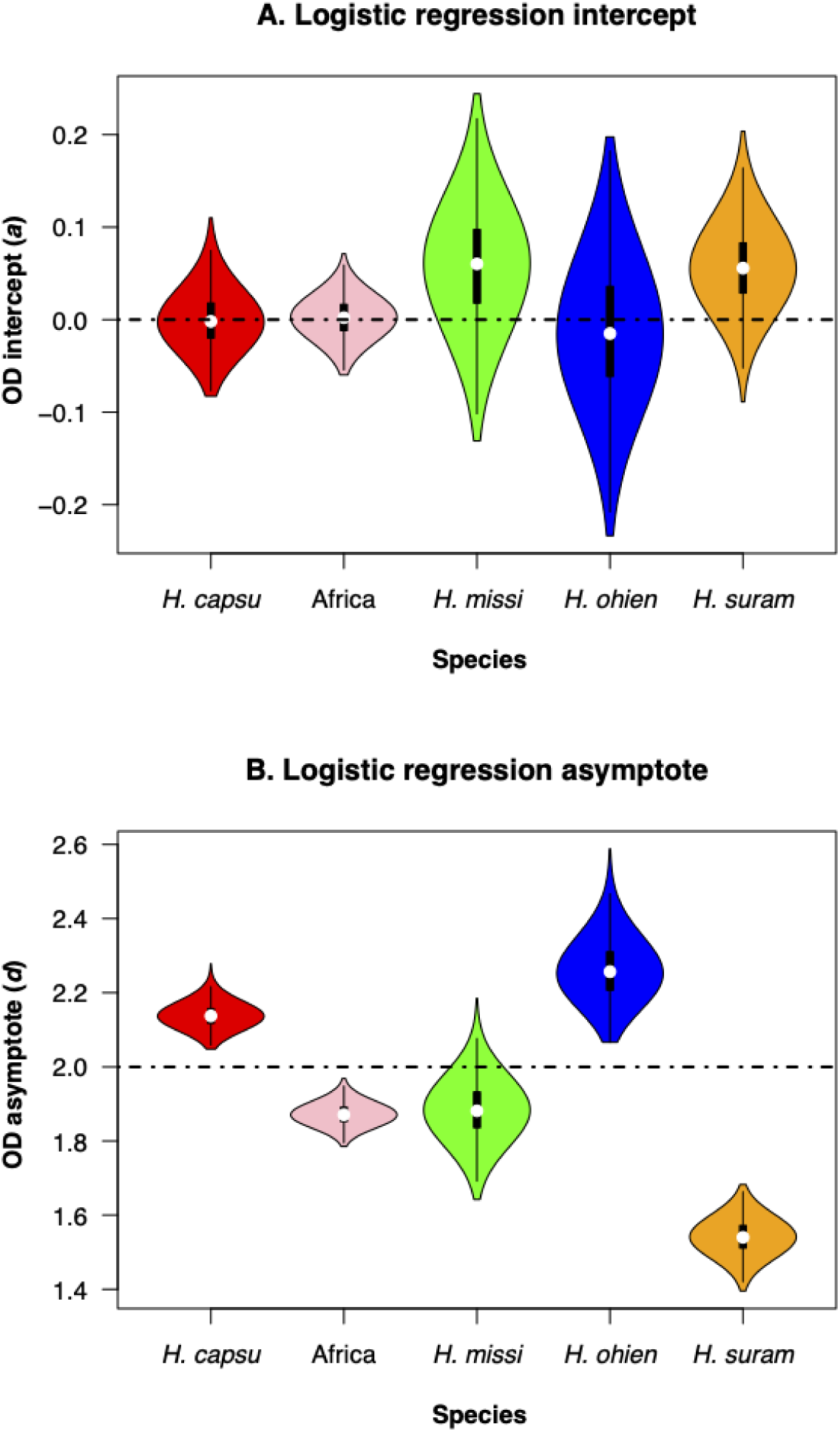
Intercept and asymptote distributions of bootstrapped regressions. **A.** Intercept (*a* in Equation 1). **B.** Asymptote (*d* in Equation 1). Each boxplot shows 1,000 values of bootstrapped non-linear regressions (shown as semitransparent lines in Figure 3). *capsu*: *H. capsulatum* ss, *H. missi*: *H. mississippiense*, *H. ohien*: *H. ohiense*, *H. suram: H. suramericanum*.

**TABLE 2.** Pairwise comparisons between two of the parameters of the logistic regression, *a* and *d*. *a* corresponds to the intercept, *d* corresponds to the asymptote. The lower triangular matrix shows the *W* from the Wilcoxon test. Upper triangular matrix shows the *p*-value. Each parameter estimate is estimated from the non-linear regression; the standard error (SE) was calculated from the distribution of the 1,000 bootstrap samplings shown in Figure 3. *capsu*: *H. capsulatum* ss; Africa: African lineage; *miss*: *H. missisippiense*; *ohien*: *H. ohiense*; *suram*: *H. suramericanum*.

| Parameter | Species | Estimate | SE<br>(bootstrap) | Species |  |  |  |  |
| --- | --- | --- | --- | --- | --- | --- | --- | --- |
|  |  |  |  | *capsu* | *africa* | *missi* | *ohien* | *suram* |
| ***a*** | *capsu* | 9.224×10<sup>-5</sup> | 9.312×10<sup>-4</sup> | * | 5.338× 10<sup>-4</sup> | 5.68× 10<sup>-10</sup> | <1 × 10<sup>-10</sup> | <1 × 10<sup>-10</sup> |
|  | *Africa* | 2.906×10<sup>-3</sup> | 6.539×10<sup>-4</sup> | 454,346 | * | 4.775× 10<sup>-3</sup> | <1 × 10<sup>-10</sup> | <1 × 10<sup>-10</sup> |
|  | *missi* | 5.708 ×10<sup>-2</sup> | 1.903×10<sup>-3</sup> | 419,071 | 462,616 | * | 3.57× 10<sup>-7</sup> | <1 × 10<sup>-10</sup> |
|  | *ohien* | -1.128×10<sup>-2</sup> | 2.272×10<sup>-3</sup> | 296,769 | 370,934 | 433,831 | * | <1 × 10<sup>-10</sup> |
|  | *suram* | 5.461×10<sup>-2</sup> | 1.294×10<sup>-4</sup> | 130,447 | 216,910 | 292,041 | 296,485 | * |
| ***d*** | *capsu* | 2.138 | 9.654×10<sup>-4</sup> | * | <1 × 10<sup>-10</sup> | <1 × 10<sup>-10</sup> | <1 × 10<sup>-10</sup> | <1 × 10<sup>-10</sup> |
|  | *africa* | 1.871 | 8.610×10<sup>-4</sup> | 998,001 | * | <1 × 10<sup>-10</sup> | <1 × 10<sup>-10</sup> | <1 × 10<sup>-10</sup> |
|  | *missi* | 1.879 | 2.318×10<sup>-3</sup> | 998,001 | 997,996 | * | <1 × 10<sup>-10</sup> | <1 × 10<sup>-10</sup> |
|  | *ohien* | 2.261 | 2.390×10<sup>-3</sup> | 993,006 | 296,455 | 2 | * | <1 × 10<sup>-10</sup> |
|  | *suram* | 1.541 | 1.427×10<sup>-3</sup> | 998,001 | 998,001 | 996,801 | 998,001 | * |

### Yeast cell size differs among *Histoplasma* species

We fit a linear model to compare yeast cell size across species. We found cell size variation among species (*F*_4,126_ = 10.31, *p* = 2.96 ×10^-7^). Table 3 shows all the pairwise comparisons among species. *Histoplasma ohiense* had a smaller size than the other four species (Table 3). The significance of the *H. ohiense*-*H. suramericanum* pairwise comparison was borderline. *Histoplasma suramericanum* had a slightly larger cell size than *H. missisippiense*, but the distributions of the two species were largely overlapping (Table 3). These results indicate that yeast cell size might serve as a diagnostic trait for *H. ohiense*, but not for the remaining *Histoplasma* species.

**TABLE 3.** Tukey HSD pairwise comparisons show that *H. ohiense* has a smaller yeast cell size than other species of *Histoplasma*. sd: Standard deviation. *capsu*: *H. capsulatum* ss; Africa: African lineage; *miss*: *H. mississippiense*; *ohien*: *H. ohiense*; *suram*: *H. suramericanum*.

|  |  |  | Species |  |  |  |  |
| --- | --- | --- | --- | --- | --- | --- | --- |
| Species | Mean (µm<sup>2</sup>) | sd | *capsu* | *africa* | *missi* | *ohii* | *suram* |
| *capsu* | 6.819 | 1.591 | * | * | * | * | * |
| *africa* | 6.511 | 1.473 | 0.949 | * | * | * | * |
| *missi* | 7.256 | 1.568 | 0.837 | 0.335 | * | * | * |
| *ohii* | 5.106 | 1.130 | 2.413×10<sup>-4</sup> | 1.768×10<sup>-3</sup> | 3.00×10<sup>-7</sup> | * | * |
| *suram* | 6.080 | 1.362 | 0.405 | 0.814 | 0.030 | 0.065 | * |

### Taxonomy

Table 4 summarizes the results of our phenotypic surveys. The combination of these traits is sufficient to differentiate between the five cryptic species of *Histoplasma*. Using these phenotyping surveys, we re-describe the three named species of *Histoplasma* and provide a dichotomous key to differentiate between the five phylogenetic species in this report.

**TABLE 4.**
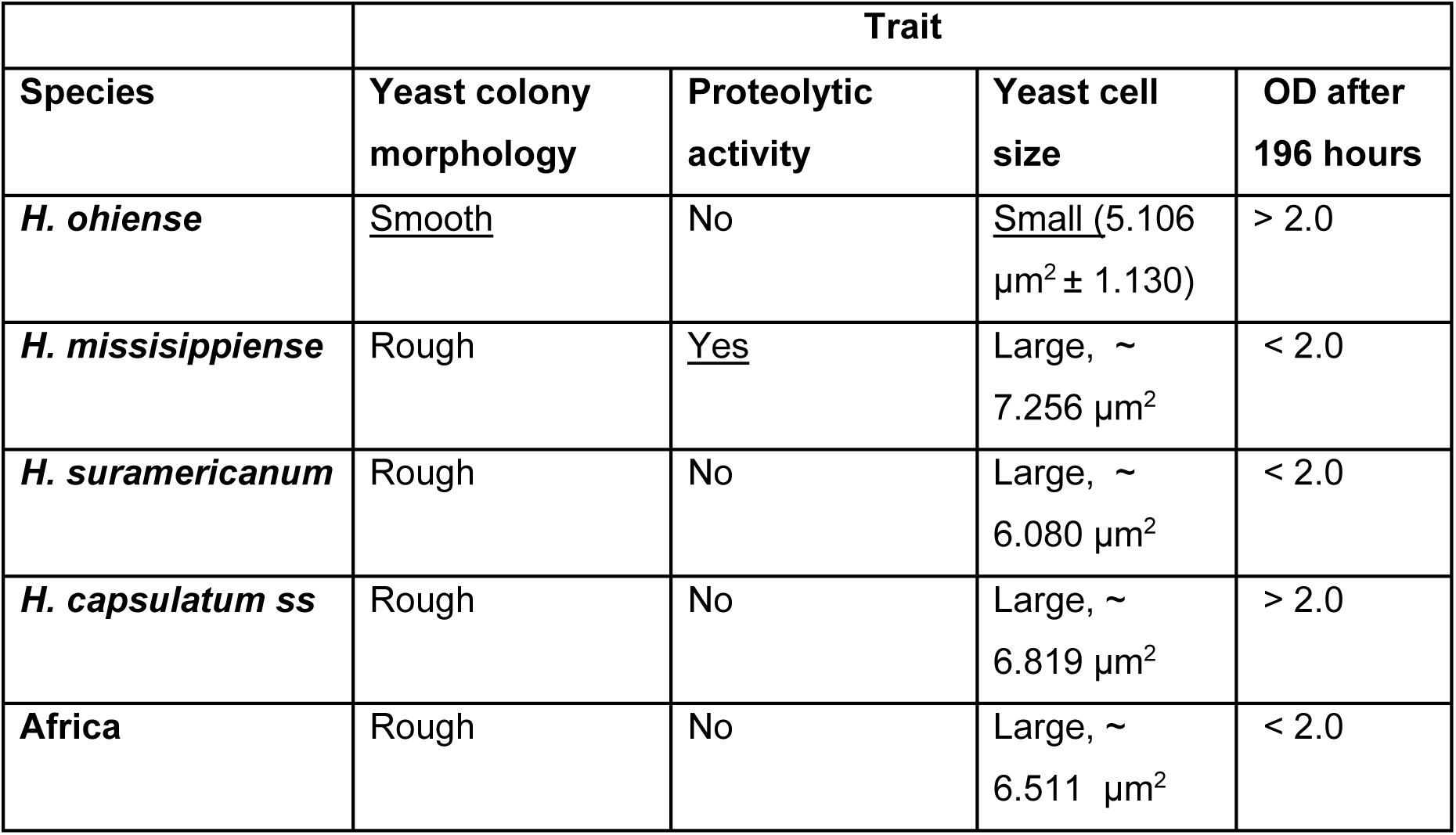
Phenotypic differences among five different species of *Histoplasma*. Species diagnostic traits are underlined.

|  | Trait |  |  |  |
| --- | --- | --- | --- | --- |
| Species | Yeast colony morphology | Proteolytic activity | Yeast cell size | OD after 196 hours |
| *H. ohii* | <u>Smooth</u> | No | <u>Small</u> (5.106 µm<sup>2</sup> ± 1.130) | > 2.0 |
| *H. mississippiense* | Rough | <u>Yes</u> | Large, ~ 7.256 µm<sup>2</sup> | < 2.0 |
| *H. suramericanum* | Rough | No | Large, ~ 6.080 µm<sup>2</sup> | < 2.0 |
| *H. capsulatum* ss | Rough | No | Large, ~ 6.819 µm<sup>2</sup> | > 2.0 |
| *Africa* | Rough | No | Large, ~ 6.511 µm<sup>2</sup> | < 2.0 |

*Histoplasma mississippiense* V.E. Sepúlveda, R. Márquez, Turissini, W.E. Goldman & Matute, sp. nov. MB XXXXXX

For a detailed description see Sepúlveda et al., mBio 8 (6): e01339-17, 12 (2017). Holotype: ATCC 38904, CBS145497 preserved in a metabolically inactive state. previously published as *Histoplasma mississippiense* V.E. Sepúlveda, R. Márquez, Turissini, W.E. Goldman & Matute, mBio 8 (6): e01339-17, 12 (2017), nom. inval., Art. 40.7 (Shenzhen), MB 823360]

*Histoplasma ohiense* V.E. Sepúlveda, R. Márquez, Turissini, W.E. Goldman & Matute, sp. nov. MB XXXXX

For a detailed description see Sepúlveda et al., mBio 8 (6): e01339-17, 13 (2017). Holotype: ATCC 26032, CBS145495 preserved in a metabolically inactive state. previously published as *Histoplasma ohiense* V.E. Sepúlveda, R. Márquez, Turissini, W.E. Goldman & Matute, mBio 8 (6): e01339-17, 12 (2017), nom. inval., Art. 40.7 (Shenzhen), MB 823360]

*Histoplasma suramericanum* V.E. Sepúlveda, R. Márquez, Turissini, W.E. Goldman & Matute, sp. nov. MB XXXXX

For a detailed description see Sepúlveda et al., mBio 8 (6): e01339-17, 13 (2017). Holotype: 3_11G, CBS145499 preserved in a metabolically inactive state. previously published as *Histoplasma suramericanum* V.E. Sepúlveda, R. Márquez, Turissini, W.E. Goldman & Matute, mBio 8 (6): e01339-17, 12 (2017), nom. inval., Art. 40.7 (Shenzhen), MB 823360]

### *Histoplasma* dichotomous key

**1A.** OD600 after 196 in HMM is higher than 2 … **2**

**1B.** OD600after 196 in HMM is lower than 2 … **3**

**2A.** Yeast colony morphology at 37°C is smooth. Yeast size area is 5.106μ^2^ ± 1.130 … *H. ohiense*

**2B**. Yeast colony morphology at 37°C is rough. Yeast size area is larger than 6.0μ^2^ … *H. capsulatum*

**3A.** Yeast colonies show proteolytic activity in HMM media at 37°C… *H. mississippiense*

**3B.** Yeast colonies show no proteolytic activity in HMM media at 37°C… **4**

**4A.** Isolate collected in the Americas. *H-antigen* precursor amplified with the forward primer 5′-CGCAGTCACCTCCATACTATC 3′ and reverse primer 5′-GCGCCGACATTAACCC-3′ (28) harbors three diagnostic SNPs (positions 591, 622, and 716, (Kasuga *et al*. 2003)). … *H. suramericanum*

**4B.** Isolate collected in Africa … African lineage

## DISCUSSION

Identifying species boundaries has been a challenge in microbial eukaryotes because producing sexual stages and making direct measurements of reproductive isolation, the signature of speciation, is usually impractical and often unfeasible (reviewed in (Taylor *et al*. 2000; Schön *et al*. 2009; Cai *et al*. 2011; Chethana *et al*. 2021)). Measuring the extent of genetic divergence, and identifying reductions in gene flow, has been a powerful substitute to uncover cryptic speciation in fungal pathogens (Birky 2013; Galtier 2019). The incorporation of genomics has opened the door to describe the evolutionary processes that govern speciation and trait diversification in fungal pathogens (Taylor *et al*. 2000; Matute and Sepúlveda 2019). Nonetheless, genome sequencing alone might be impractical for the identification of pathogens, particularly in clinical settings. In this study, we report phenotypic differences that are sufficient to identify three of five named species of *Histoplasma*, and revise their taxonomic status. In particular, we report that *H. ohiense* can be identified by its characteristic smooth colonies and small cell size, and *H. mississippiense* by its extracellular proteolytic activity. *Histoplasma suramericanum*, *H. mississippiense*, and the African lineage all have an OD600 asymptote at 196 hours lower than 2, but only *H. mississippiense has* extracellular proteolytic activity. This makes *H. suramericanum* and the African lineage the only species that cannot be differentiated by the morphological traits that we describe in this survey. Nonetheless, *H. suramericanum* seems to be restricted to the American continent, and the African lineage to Africa. Additionally, the two species can be discriminated with PCR probes (Kasuga *et al*. 1999, 2003; Sepúlveda *et al*. 2017). Our results are of importance to evolutionary and clinical mycologists alike because the diagnosis of species boundaries is the first step to understanding evolutionary dynamics, broadly defined, and could shed light into the evolution of different virulence mechanisms.

Other studies have reported differences in the morphology of *Histoplasma* isolates, and among clusters of genotypes. Okeke and Muller (Okeke and Müller 1991) described the presence of extracellular collagenolytic proteinases by *Histoplasma capsulatum* var. *duboisii* and *Histoplasma capsulatum* var. *capsulatum*. Since these classifications do not follow a phylogenetic framework (Kasuga *et al*. 1999, 2003), the results are not immediately comparable. Importantly, our results are consistent with Zarnowski *et al*. (2007), where the extracellularly-secreted serine protease activity was restricted to *H. mississippiense* isolates (formerly known as RFLP class 1 or NAm 1 clade, (Keath *et al*. 1992; Kasuga *et al*. 1999, 2003)), demonstrating the methodology and media used in both studies is suitable for the selective identification of *H. mississippiense*, and can be used reliably. To date, the role of the extracellularly-secreted serine protease activity in *H. mississippiense* virulence remains unexplored. Muotoe-Okafor et al. (Muotoe-Okafor *et al*. 1996) detected a similar proteolytic activity in a cluster of African samples. Since no other species besides the Africa clade has been isolated from Africa, these results seem to indicate that some African isolates, but not the ones included in this study, might have serine protease activities similar to the ones in *H. mississippiense*. These two species are not sisters in the phylogenetic tree and thus, these results suggest that this *H. mississippiense* specific proteolytic activity might be explained by genomic changes that are specific to that lineage, and might have also evolved in the African clade through parallel mutation or introgression. A second, arguably less likely, possibility is that other species of *Histoplasma* (*H. ohiense*, *H. capsulatum* ss) lost the serine proteinase activity independently. If that is the case, these two species should harbor serine proteinase pseudogenes. Now that species boundaries have been identified in *Histoplasma*, studies dissecting the processes that lead to serine proteinases in this genus of fungi are within reach.

Yeast colony morphology is arguably the most systematically studied phenotypic difference in *Histoplasma*. The existence of smooth and rough colony morphology in *Histoplasma* was first reported as early as 1987 (Klimpel and Goldman 1987). Genetic analyses suggested that the smooth phenotype was exclusive to a cluster of genotypes (RFLP2, *NAm* 2), now dubbed *H. ohiense*. Detailed studies of the cell wall with transmission electron microscopy demonstrated that reference strains of *H. ohiense* and *H. capsulatum* differ in their cell wall thickness, with *H. capsulatum* yeast cells showing a greater cell wall thickness compared to *H. ohiense* and that *AGS1* expression is dispensable for *H. ohiense* virulence (Edwards et al, 2011). α-(1, 3)-glucan is required for virulence in *H. capsulatum* and *H. mississippiense* (Rappleye et al, 2004 and Sepulveda et al, 2014 respectively); smooth mutants become avirulent once they are unable to produce this polysaccharide. *Histoplasma ohiense* has smooth colonies and lacks α-(1, 3)-glucan, and yet it remains virulent. The dissection of the genetic basis of differences in virulence between *Histoplasma* species is a prime example of the importance of understanding species boundaries in eukaryotic pathogens.

There is extensive precedent that once fungal species are identified, phenotypic differences between the newfound taxa are subsequently found. In the case of *Coccidioides—*the first fungal pathogen to undergo a taxonomic revision (Koufopanou *et al*. 1997, 2001; Kirkland and Fierer 2018), the two different species, *C. posadassi* and *C. immitis* show differences in thermotolerance, which might be of importance for yeast-to-mycelium transformation and in determining their geographic range (Mead *et al*. 2020). Similarly, different species of *Paracoccidioides* show differences in antifungal resistance (Cruz *et al*. 2013) and yeast morphology (Turissini *et al*. 2017), but also in the host response they elicit in their mammalian hosts (Teixeira *et al*. 2014; Siqueira *et al*. 2015; De Macedo *et al*. 2019; Hahn *et al*. 2019). Even though reports of phenotypic variability existed in these fungi (e.g., (Klimpel and Goldman 1988)), ascribing these differences to species boundaries was only possible once isolated lineages were described in genera that were considered monotypic for almost 100 years.

Our study focuses on five lineages identified through genome sequencing, but there is precedent suggesting that *Histoplasma* contains additional differentiated lineages. Initial surveys using multilocus-sequence typing reported the existence of over a dozen lineages that might fulfill the criteria for phylogenetic species. Sequencing of samples from other locations has revealed additional clades that fulfill the requirements to be considered monophyletic species (Rio de Janeiro in Brazil: (Almeida-Silva *et al*. 2021; India: Jofre *et al*. 2022). Genomic studies that quantified the different trajectories along the genome in a worldwide sample are sorely needed. Multiple studies have reported inter-isolate differences in the *Histoplasma* genus, but a systematic survey that includes not only reference isolates, but also a variety of other strains is needed. For example, the reference isolate of *H. ohiense* (G217B) is more virulent than its counterpart in *H. mississippiense* (WU24) in mouse inoculations (Sepúlveda *et al*. 2014). Similarly, a clinical isolate of *H. mississippiense* is more resistant to fluconazole than the reference isolate of *H. ohiense* (Goughenour *et al*. 2015; Goughenour and Rappleye 2017), highlighting the importance of considering which species is responsible for causing disease in a patient when deciding on the course of treatment. Finally, the reference strain of *H. capsulatum* (G186A) induces a higher infiltration of monocytic cells in the lungs of mice inoculated with a low dose (10^3^ yeast) than the representative isolates of *H. mississippiense* and *H. ohiense* (Sepúlveda *et al*. 2014); Jones et al. 2021). All these surveys suffer from the same shortcoming, which is that differences between isolates might not be representative of the differences among species. Nonetheless, they are powerful starting points to propel surveys that quantify the extent of inter- and intraspecies variation.

The case of *Histoplasma* will require a more systematic exploration than that of *Coccidioides* or *Paracoccidioides* because the number of lineages in *Histoplasma* appears to be much higher than in either of those other fungal pathogens. There is already indication that other unnamed *Histoplasma* lineages show important phenotypic differences. For example, a phylogenetic species restricted to Rio de Janeiro, Brazil seems to have a higher likelihood of causing hemorrhages than other genotypes (Almeida-Silva *et al*. 2021). It is imperative as we define species boundaries that we also make a systematic effort to find phenotypic traits to aid species identification, as they can become useful tools in the clinical setting and could have an impact on the type of antifungal therapy used to treat infections. Our work demonstrates that morphological differences among *Histoplasma* species do exist and provides a blueprint for future surveys.

## ACKNOWLEDGEMENTS

We would like to thank our reviewers and members of the Matute lab for helpful comments. This work was supported by the National Institute of General Medical Sciences of the National Institutes of Health (NIH) under Award R01AI153523 to DRM. The funders had no role in study design, data collection and analysis, decision to publish, or preparation of the manuscript.

## SUPPLEMENTARY MATERIAL

**TABLE S1.** AIC values for the best fitting dose-response and linear functions for the growth curve of each *Histoplasma* species.

|  | AIC |  |
| --- | --- | --- |
| Species | Linear | Dose-response |
| *H. suramericanum* | -5.038041 | -20.2201 |
| *H. ohiense* | 16.69309 | -8.40372 |
| *H. mississippiense* | 9.61138 | -8.56707 |
| *H. capsulatum* ss | 13.66652 | -41.2338 |
| Africa | 1.258701 | -71.5962 |

**TABLE S2.**
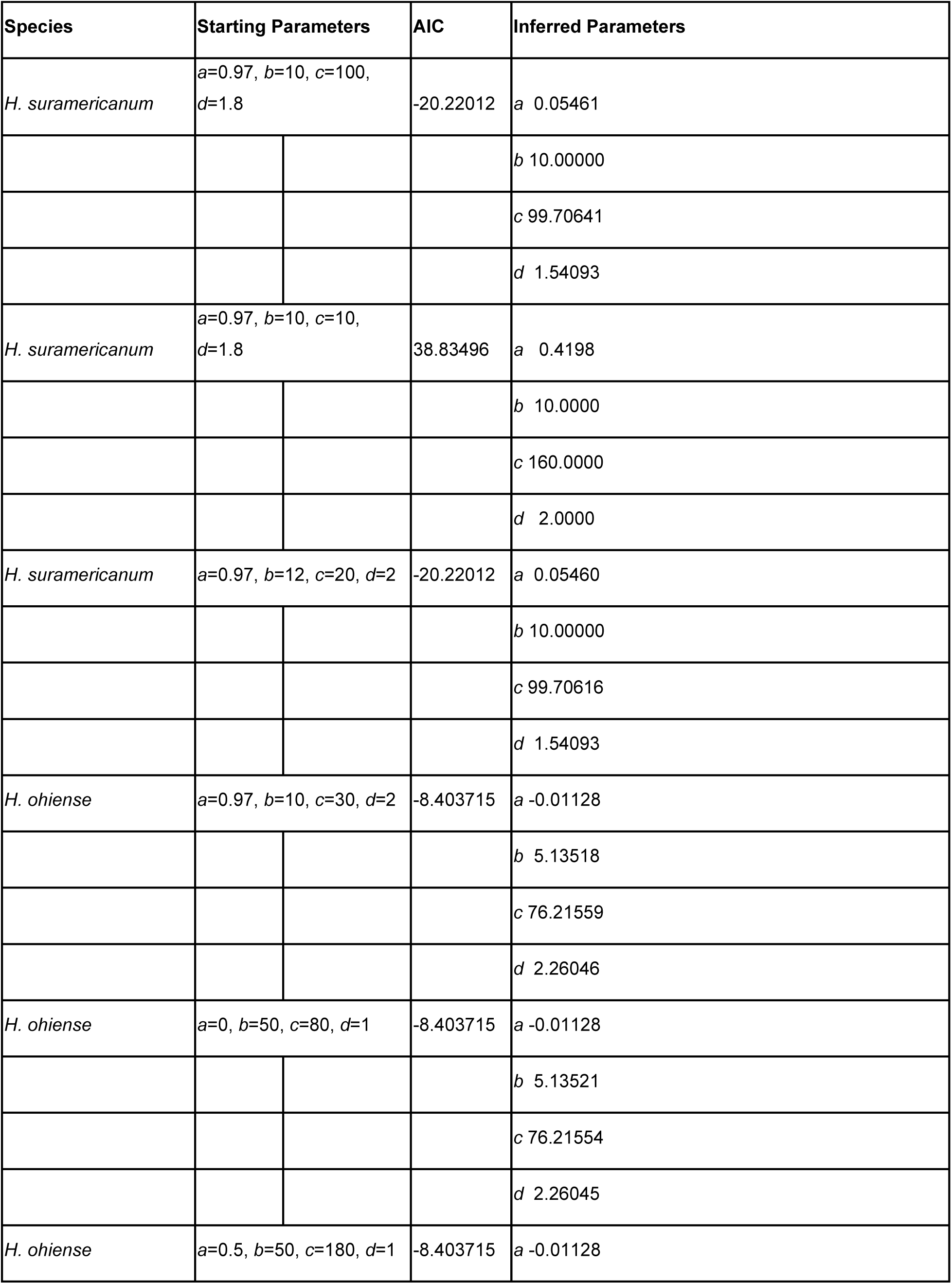
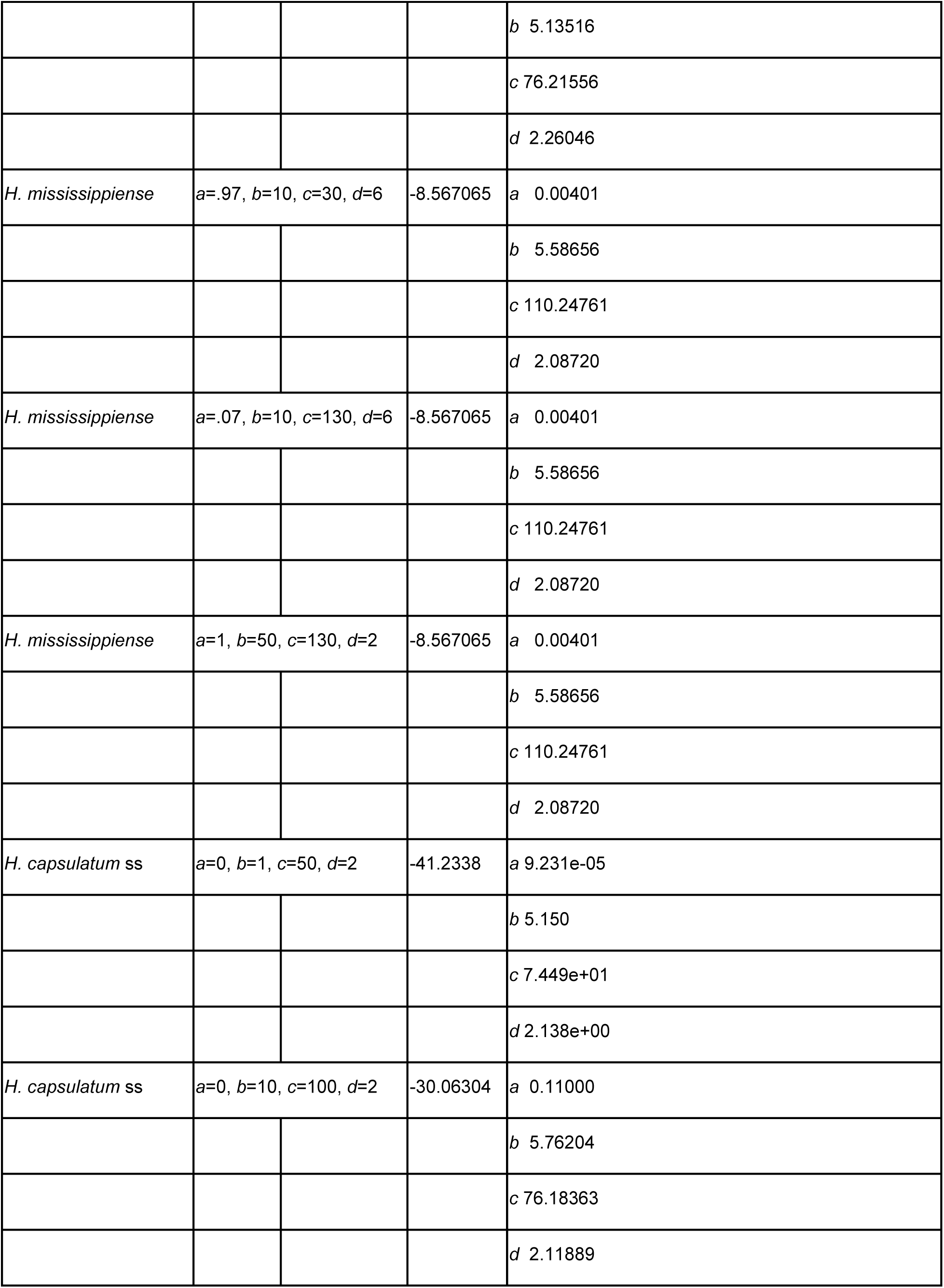
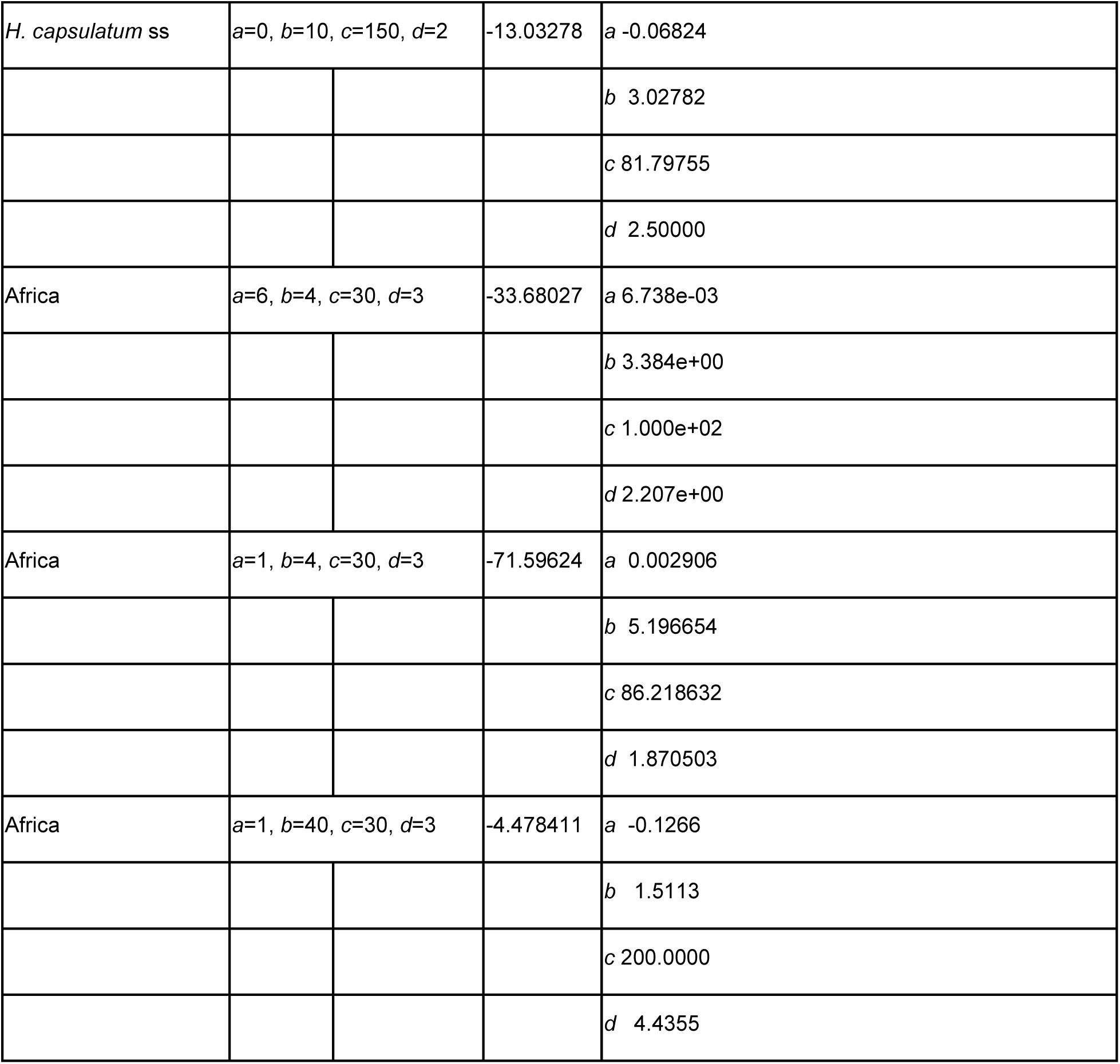
**Effect of different optimization values for the dose-response functions.**

## REFERENCES

Adderson E. E., 2004 Histoplasmosis in a pediatric oncology center. The Journal of pediatrics 144: 100–106.

Adenis A. A., A. Valdes, C. Cropet, O. Z. McCotter, G. Derado, et al., 2018 Burden of HIV-associated histoplasmosis compared with tuberculosis in Latin America: a modelling study. Lancet Infect Dis 18: 1150–1159. 10.1016/S1473-3099(18)30354-2

Ajello L., 1968 Comparative Morphology and Immunology of Members of the Genus *Histoplasma*: A Review. Mycoses 11: 507–514. 10.1111/j.1439-0507.1968.tb03370.x

Akaike H., 1973 Maximum likelihood identification of Gaussian autoregressive moving average models. Biometrika 60: 255–265.

Almeida-Silva F., M. de Melo Teixeira, D. R. Matute, M. de Faria Ferreira, B. M. Barker, et al., 2021 Genomic Diversity Analysis Reveals a Strong Population Structure in *Histoplasma capsulatum* LAmA (*Histoplasma suramericanum*). J Fungi (Basel) 7: 865. 10.3390/jof7100865

Armstrong P. A., B. R. Jackson, D. Haselow, V. Fields, M. Ireland, et al., 2018 Multistate epidemiology of histoplasmosis, United States, 2011–20141. Emerging infectious diseases 24: 425.

Baty F., M.-L. Delignette-Muller, S. Charles, J.-P. Flandrois, C. Ritz, et al., 2013 Package ‘nlstools.’ R Foundation for Statistical Computing, Vienna.

Baty F., C. Ritz, S. Charles, M. Brutsche, J.-P. Flandrois, et al., 2015 A toolbox for nonlinear regression in R: the package *nlstools*. Journal of Statistical Software 66: 1–21.

Benedict K., and R. K. Mody, 2016 Epidemiology of histoplasmosis outbreaks, United States, 1938–2013. Emerging infectious diseases 22: 370.

Birky C. W., 2013 Species detection and identification in sexual organisms using population genetic theory and DNA sequences. PLoS One 8: e52544. 10.1371/journal.pone.0052544

Bongomin F., R. Kwizera, and D. W. Denning, 2019 Getting Histoplasmosis on the Map of International Recommendations for Patients with Advanced HIV Disease. J Fungi (Basel) 5: 80. 10.3390/jof5030080

Cai L., T. Giraud, N. Zhang, D. Begerow, G. Cai, et al., 2011 The evolution of species concepts and species recognition criteria in plant pathogenic fungi. Fungal Diversity 50: 121–133.

Cano M. V., and R. A. Hajjeh, 2001 The epidemiology of histoplasmosis: a review., pp. 109–118 in Seminars in respiratory infections,.

Chethana K. T., I. S. Manawasinghe, V. G. Hurdeal, C. S. Bhunjun, M. A. Appadoo, et al., 2021 What are fungal species and how to delineate them? Fungal Diversity 109: 1–25.

Cruz R. C., S. M. C. Werneck, C. S. Oliveira, P. C. Santos, B. M. Soares, et al., 2013 Influence of Different Media, Incubation Times, and Temperatures for Determining the MICs of Seven Antifungal Agents against *Paracoccidioides brasiliensis* by Microdilution. J Clin Microbiol 51: 436–443. 10.1128/JCM.02231-12

De Macedo P. M., M. de M. Teixeira, B. M. Barker, R. M. Zancope-Oliveira, R. Almeida-Paes, et al., 2019 Clinical features and genetic background of the sympatric species *Paracoccidioides brasiliensis* and *Paracoccidioides americana*. PLoS neglected tropical diseases 13: e0007309.

Domer J. E., 1971 Monosaccharide and Chitin Content of Cell Walls of *Histoplasma capsulatum* and *Blastomyces dermatitidis*. J Bacteriol 107: 870–877. 10.1128/jb.107.3.870-877.1971

Edwards J. A., E. A. Alore, and C. A. Rappleye, 2011 The Yeast-Phase Virulence Requirement for α-Glucan Synthase Differs among *Histoplasma capsulatum* Chemotypes. Eukaryot Cell 10: 87–97. 10.1128/EC.00214-10

Freifeld A. G., P. C. Iwen, B. L. Lesiak, R. K. Gilroy, R. B. Stevens, et al., 2005 Histoplasmosis in solid organ transplant recipients at a large Midwestern university transplant center. Transplant Infectious Dis 7: 109–115. 10.1111/j.1467-8365.2005.00105.x

Galtier N., 2019 Delineating species in the speciation continuum: A proposal. Evolutionary applications 12: 657–663.

Goughenour K. D., J.-M. Balada-Llasat, and C. A. Rappleye, 2015 Quantitative Microplate-Based Growth Assay for Determination of Antifungal Susceptibility of *Histoplasma capsulatum* Yeasts, (D. W. Warnock, Ed.). J Clin Microbiol 53: 3286–3295. 10.1128/JCM.00795-15

Goughenour K. D., and C. A. Rappleye, 2017 Antifungal therapeutics for dimorphic fungal pathogens. Virulence 8: 211–221. 10.1080/21505594.2016.1235653

Hage C., M. Azar, N. Bahr, J. Loyd, and L. Wheat, 2015 Histoplasmosis: Up-to-Date Evidence-Based Approach to Diagnosis and Management. Semin Respir Crit Care Med 36: 729– 745. 10.1055/s-0035-1562899

Hahn R. C., A. M. Rodrigues, P. P. Della Terra, A. F. Nery, H. D. Hoffmann-Santos, et al., 2019 Clinical and epidemiological features of paracoccidioidomycosis due to *Paracoccidioides lutzii*. PLoS neglected tropical diseases 13: e0007437.

Hess J., A. Fondell, N. Fustino, J. Malik, and C. Rokes, 2017 Presentation and treatment of histoplasmosis in pediatric oncology patients: case series and review of the literature. Journal of Pediatric Hematology/Oncology 39: 137–140.

Jofre G. I., A. Singh, H. Mavengere, G. Sundar, E. D’Agostino, et al., 2022 An Indian lineage of *Histoplasma* with strong signatures of differentiation and selection. Fungal Genet Biol 158: 103654. 10.1016/j.fgb.2021.103654

Jones GS, Sepúlveda VE, Goldman WE. 2020. Biodiverse *Histoplasma* Species Elicit Distinct Patterns of Pulmonary Inflammation following Sublethal Infection. mSphere. 5:e00742–20. doi: 10.1128/mSphere.00742-20.

Kasuga T., J. W. Taylor, and T. J. White, 1999 Phylogenetic relationships of varieties and geographical groups of the human pathogenic fungus *Histoplasma capsulatum* Darling. J Clin Microbiol 37: 653–663. 10.1128/JCM.37.3.653-663.1999

Kasuga T., T. J. White, G. Koenig, J. McEwen, A. Restrepo, et al., 2003 Phylogeography of the fungal pathogen *Histoplasma capsulatum*. Mol Ecol 12: 3383–3401. 10.1046/j.1365-294x.2003.01995.x

Kauffman C. A., 2007 Histoplasmosis: a Clinical and Laboratory Update. Clin Microbiol Rev 20: 115–132. 10.1128/CMR.00027-06

Keath E. J., G. S. Kobayashi, and G. Medoff, 1992 Typing of *Histoplasma capsulatum* by restriction fragment length polymorphisms in a nuclear gene. Journal of clinical microbiology 30: 2104–2107.

Kirkland T. N., and J. Fierer, 2018 *Coccidioides immitis* and *posadasii*; A review of their biology, genomics, pathogenesis, and host immunity. Virulence 9: 1426–1435. 10.1080/21505594.2018.1509667

Klimpel K. R., and W. E. Goldman, 1987 Isolation and characterization of spontaneous avirulent variants of *Histoplasma capsulatum*. Infect Immun 55: 528–533. 10.1128/iai.55.3.528-533.1987

Klimpel K. R., and W. E. Goldman, 1988 Cell walls from avirulent variants of *Histoplasma capsulatum* lack alpha-(1,3)-glucan. Infect Immun 56: 2997–3000. 10.1128/iai.56.11.2997-3000.1988

Koufopanou V., A. Burt, and J. W. Taylor, 1997 Concordance of gene genealogies reveals reproductive isolation in the pathogenic fungus *Coccidioides immitis*. Proc Natl Acad Sci U S A 94: 5478–5482. 10.1073/pnas.94.10.5478

Koufopanou V., A. Burt, T. Szaro, and J. W. Taylor, 2001 Gene genealogies, cryptic species, and molecular evolution in the human pathogen *Coccidioides immitis* and relatives (Ascomycota, Onygenales). Mol Biol Evol 18: 1246–1258. 10.1093/oxfordjournals.molbev.a003910

Kwon-Chung K. J., 1972a Sexual Stage of *Histoplasma capsulatum*. Science 175: 326–326. 10.1126/science.175.4019.326

Kwon-Chung K. J., 1972b *Emmonsiella capsulata* : Perfect State of *Histoplasma capsulatum*. Science 177: 368–369. 10.1126/science.177.4046.368.b

Kwon-Chung K. J., 1973 Studies on *Emmonsiella capsulata* I. Heterothallism and development of the ascocarp. Mycologia 109–121.

Manos N. E., S. H. Ferebee, and W. F. Kerschbaum, 1956 Geographic variation in the prevalence of histoplasmin sensitivity. Diseases of the chest 29: 649–668.

Matute D. R., and V. E. Sepúlveda, 2019 Fungal species boundaries in the genomics era. Fungal Genet Biol 131: 103249. 10.1016/j.fgb.2019.103249

Maxwell C. S., V. E. Sepulveda, D. A. Turissini, W. E. Goldman, and D. R. Matute, 2018 Recent admixture between species of the fungal pathogen *Histoplasma*. Evolution Letters 2: 210–220.

Mead H. L., P. S. Hamm, I. N. Shaffer, M. de M. Teixeira, C. S. Wendel, et al., 2020 Differential thermotolerance adaptation between species of *Coccidioides*. Journal of Fungi 6: 366.

Muniz M. M., C. N. Sousa, M. M. E. Oliveira, C. V. Pizzini, M. A. Almeida, et al., 2014 Sexual variability in *Histoplasma capsulatum* and its possible distribution: what is going on? Revista Iberoamericana De Micologia 31: 7–10.

Muotoe-Okafor F. A., H. C. Gugnani, and O. O. Obidoa, 1996 Extracellular proteolytic enzyme activity of *Histoplasma capsulatum* var.*duboisii*. Mycopathologia 133: 129–133. 10.1007/BF02373018

Myint T., N. Leedy, E. Villacorta Cari, and L. J. Wheat, 2020 HIV-Associated Histoplasmosis: Current Perspectives. HIV Volume 12: 113–125. 10.2147/HIV.S185631

Okeke C. N., and J. Müller, 1991 In vitro production of extracellular elastolytic proteinase by *Histoplasma capsulatum* var. *duboisii* and *Histoplasma capsulatum* var. *capsulatum* in the yeast phase: Die Bildung extrazellulärer elastolytischer Proteinase in vitro durch *Histoplasma capsulatum* var. *duboisii* und *Histoplasma capsulatum* var. *capsulatum* in der Hefephase. Mycoses 34: 461–467. 10.1111/j.1439-0507.1991.tb00861.x

Oladele R. O., O. O. Ayanlowo, M. D. Richardson, and D. W. Denning, 2018 Histoplasmosis in Africa: an emerging or a neglected disease? PLoS neglected tropical diseases 12: e0006046.

R Core Team Rf., 2018 R: A language and environment for statistical computing Rappleye C. A., J. T. Engle, and W. E. Goldman, 2004 RNA interference in *Histoplasma capsulatum* demonstrates a role for α-(1,3)-glucan in virulence. Molecular Microbiology 53: 153–165. 10.1111/j.1365-2958.2004.04131.x

Reiss E., 1977 Serial enzymatic hydrolysis of cell walls of two serotypes of yeast-form *Histoplasma capsulatum* with alpha(1 leads to 3)-glucanase, beta(1 leads to 3)-glucanase, pronase, and chitinase. Infect Immun 16: 181–188. 10.1128/iai.16.1.181-188.1977

Reiss E., S. E. Miller, W. Kaplan, and L. Kaufman, 1977 Antigenic, Chemical, and Structural Properties of Cell Walls of *Histoplasma capsulatum* Yeast-Form Chemotypes 1 and 2 After Serial Enzymatic Hydrolysis. Infect Immun 16: 690–700. 10.1128/iai.16.2.690-700.1977

Schneider C. A., W. S. Rasband, and K. W. Eliceiri, 2012 NIH Image to ImageJ: 25 years of image analysis. Nat Methods 9: 671–675. 10.1038/nmeth.2089

Schön I., K. Martens, and P. Dijk (Eds.), 2009 Lost Sex: The Evolutionary Biology of Parthenogenesis. Springer Netherlands, Dordrecht.

Scully M. C., and J. W. Baddley, 2018 Epidemiology of histoplasmosis. Current Fungal Infection Reports 12: 51–58.

Sepúlveda V. E., C. L. Williams, and W. E. Goldman, 2014 Comparison of phylogenetically distinct *Histoplasma* strains reveals evolutionarily divergent virulence strategies. mBio 5: e01376–01314. 10.1128/mBio.01376-14

Sepúlveda V. E., R. Márquez, D. A. Turissini, W. E. Goldman, and D. R. Matute, 2017 Genome sequences reveal cryptic speciation in the human pathogen *Histoplasma capsulatum*. MBio 8: e01339–17.

Siqueira I. M., C. L. F. Fraga, A. C. Amaral, A. C. O. Souza, M. S. Jerônimo, et al., 2015 Distinct patterns of yeast cell morphology and host responses induced by representative strains of *Paracoccidioides brasiliensis* (Pb18) and *Paracoccidioides lutzii* (Pb01). Sabouraudia 54: 177–188.

Taylor J. W., D. J. Jacobson, S. Kroken, T. Kasuga, D. M. Geiser, et al., 2000 Phylogenetic species recognition and species concepts in fungi. Fungal genetics and biology 31: 21– 32.

Teixeira M. de M., R. C. Theodoro, F. F. M. de Oliveira, G. C. Machado, R. C. Hahn, et al., 2014 *Paracoccidioides lutzii* sp. nov.: biological and clinical implications. Med Mycol 52: 19–28. 10.3109/13693786.2013.794311

Teixeira M. de M., J. S. L. Patané, M. L. Taylor, B. L. Gómez, R. C. Theodoro, et al., 2016 Worldwide Phylogenetic Distributions and Population Dynamics of the Genus *Histoplasma*. PLoS Negl Trop Dis 10: e0004732.

Turissini D. A., O. M. Gomez, M. M. Teixeira, J. G. McEwen, and D. R. Matute, 2017 Species boundaries in the human pathogen *Paracoccidioides*. Fungal Genet Biol 106: 9–25.

Voorhies M., S. Cohen, T. P. Shea, S. Petrus, J. F. Muñoz, et al., 2022 Chromosome-level genome assembly of a human fungal pathogen reveals synteny among geographically distinct species. MBio 13: e02574–21.

Worsham P. L., and W. E. Goldman, 1988 Quantitative plating of *Histoplasma capsulatum* without addition of conditioned medium or siderophores. Journal of Medical and Veterinary Mycology 26: 137–143.

Zarnowski R., P. A. Connolly, L. J. Wheat, and J. P. Woods, 2007 Production of extracellular proteolytic activity by *Histoplasma capsulatum* grown in *Histoplasma*-macrophage medium is limited to restriction fragment length polymorphism class 1 isolates. Diagnostic microbiology and infectious disease 59: 39–47.

